# Trust Modulates Speech Entrainment: Enhanced Cortical Tracking for Low-Trust Speakers

**DOI:** 10.64898/2026.03.11.711118

**Authors:** Jaimy A. Hannah, Giovanni M. Di Liberto

**Affiliations:** School of Computer Science and Statistics, The University of Dublin, Trinity College, Dublin, Ireland; ADAPT Centre; Trinity College Institute of Neuroscience, The University of Dublin, Trinity College, Dublin, Ireland; Trinity Centre for Biomedical Engineering, Ireland

**Keywords:** EEG, temporal response function, cortical tracking, speech entrainment, speech perception, trust

## Abstract

Trust is a critical component of human communication, providing a foundation for understanding, information exchange, and social coordination. Much of the research on trust in speech communication has focused on how vocal characteristics impact perceived trustworthiness. However, little is known about how trust in a speaker affects the neural processing of speech. Here, we demonstrate a two-stage experimental framework to study that question using non-invasive EEG. First, participants engage in a trust-building stage, where they play an investment game with fictional characters, each paired with a distinctive voice and trustworthiness level (i.e., frequency and magnitude of lies). Next, participants engage in a story-listening stage, in which they are presented with stories from the same characters. Data acquired from twenty young adults confirm a statistically significant correlation between the perceived and actual trustworthiness of the fictional characters. Cortical speech tracking was quantified using a temporal response function (TRF) analysis on the EEG data. We found that the trustworthiness established during the trust-building stage influenced the cortical tracking of speech in the subsequent story-listening stage, with lower trustworthiness corresponding to a stronger cortical tracking of speech. Interestingly, trustworthiness selectively modulated tracking strength, with no statistically significant changes in the spatial and temporal TRF dynamics.

## Introduction

Trust is a critical component of social interaction, with significant impacts on communication, collaboration, and cognition (Mayer et al., 1995; Rousseau et al., 1998). Because speech communication is one of our primary social tools, trust has direct consequences for how we listen and what we take away from an interaction. That is, if we believe someone to be trustworthy, we will receive what they say – and how they say it – differently than we would an untrustworthy communication partner.

Research on trust and speech communication has extensively investigated the specific linguistic and voice characteristics that impact perceived trustworthiness (e.g., Belin et al., 2017; Burrell et al., 2024; Elkins & Derrick, 2013; Goupil et al., 2021; Jiang et al., 2020; Levitan & Hirschberg, 2022; for a systematic review, see: Maltezou-Papastylianou et al., 2025). Trustworthiness can even be inferred from a single word (e.g., “*hello*”) via cues in voice timbre and pitch dynamics, especially the starting and ending F_0_ (Belin et al., 2017). Fluency and speech rate also correlate with perceived trustworthiness, with filler words, disfluencies (especially pauses), and variable speech rate being associated with lower trust (Burrell et al., 2024; Chen et al., 2020; Manson et al., 2013; Miller et al., 1976; Unkelbach et al., 2011).

While a large body of work has investigated how these acoustic cues relate with perceived trustworthiness, less is known about how established trust or distrust in a talker influences how their speech is received and processed by the listener’s brain. As in the classic fable “*The Boy Who Cried Wolf"*, where repeated deception erodes credibility and prompts inaction when it matters, established trustworthiness can exert a profound influence on our actions. Here, we test whether this influence extends to the neural processing of speech. To relate these two factors, trust and the neural processing of speech, we propose a novel experimental framework combining concepts from trust research and speech neurophysiology.

The proposed two-stage experimental framework (**Figure 1**) starts with a *trust-building stage* inspired by investment-based trust games from behavioural research (Berg et al., 1995; Camerer & Weigelt, 1988). In the classic paradigm, a Trustor decides how much money to give a Trustee, who receives three times as much, and then decides how much to return to the Trustor. This simple, single-trial paradigm produces numerical indices of trust-in-others for the Trustor and trustworthiness for the Trustee (Berg et al., 1995; Johnson & Mislin, 2011). In the present experiment, we adopted a multi-turn version (King-Casas et al., 2005), enabling participants (acting as Trustors) to establish trust (or lack thereof) with multiple fictional characters while hearing sufficient speech material to familiarise with the character voice. Our manipulation was designed to establish differing levels of trust through repeated interactions in which participants learned that different speakers varied in their deceptive behaviour. This approach is grounded in two commonly cited definitions of trust (Mayer et al., 1995; Rousseau et al., 1998), which define trust in terms of a willingness to be vulnerable to the actions of others based on expectations about how they will act. Participants had to decide how vulnerable to be (i.e., how many coins to invest) based on their experiences. More positive and reliable interactions should lead to greater willingness to be vulnerable, ultimately leading to higher trust.

**Figure 1.**
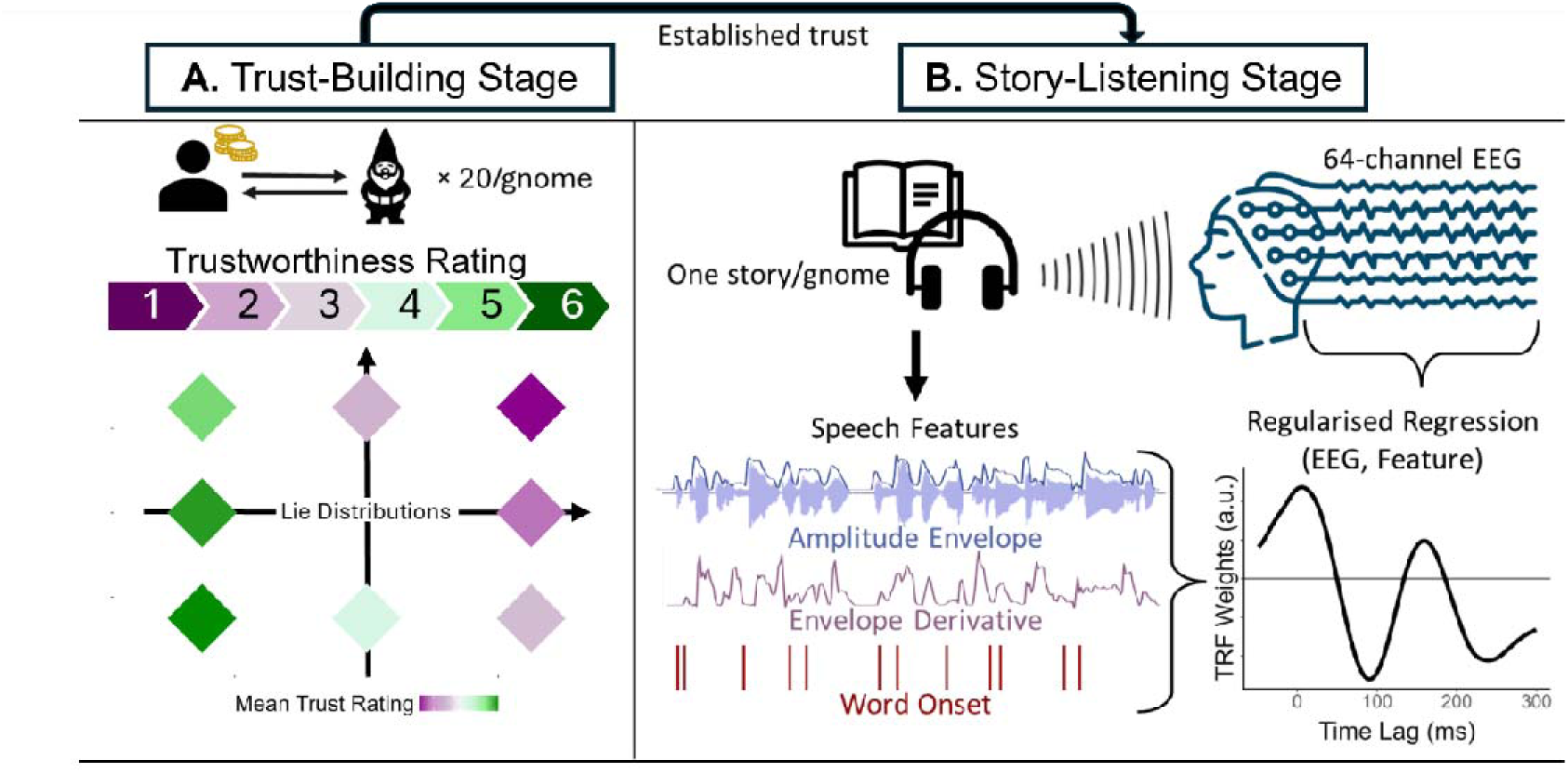
Experimental procedure and analysis method. **(A)** Participants engaged in a trust-building stage, where they played an investment-based trust game with fictional gnome characters, each associated with specific colour and voice (randomised across participants). The gnomes lied at different frequencies and with different severities as depicted by the distribution in bottom of this panel. This procedure led to establishing varying degrees of trust with the different characters (each paired with a specific voice). Participants then reported a trustworthiness rating for each character, with values ranging from 1 (not at all trustworthy) to 6 (extremely trustworthy). Participant trust ratings closely followed the distribution of lies, with the gnomes lying more often and more severely (top right) receiving the lowest trust ratings (dark purple) and the gnomes lying the least often and least severely (bottom left) receiving the highest trust ratings (dark green). **(B)** Next, participants engaged in a story-listening stage, where they were presented with stories by these same gnome characters. The central hypothesis was that established trustworthiness resulting from stage one influences the neural processing of speech in the story-listening stage. To test that hypothesis, we modelled the linear mapping of speech features (amplitude envelope, envelope derivative, and word onsets) to the corresponding EEG responses using a multivariate temporal response function (TRF) analysis, providing measurements of cortical speech tracking.

Immediately after the trust-building stage, participants engaged in the *story-listening stage*. This task has been widely studied in the speech neurophysiology literature (Brodbeck et al., 2018; Broderick et al., 2018; Crosse et al., 2021; Lalor & Foxe, 2010) and is known to elicit cortical activity that can be analysed to measure the neural processing of speech. Participants were asked to make a trust judgement after hearing each story, which would combine their (dis)trust on the fictional character and what they heard in the specific story. Here, neural activity was recorded non-invasively with high-density scalp electroencephalography (EEG) and was analysed using the temporal response function framework. We hypothesised that trust established during the *trust-building stage* would influence EEG signatures of speech in the subsequent *story-listening stage*, thereby allowing us to test how trust shapes the neural processing of speech. When listeners perceive a speaker as unreliable, they may engage in enhanced monitoring of the incoming message to detect inconsistencies or deception, requiring greater attention and increasing cognitive load. Previous work has shown that speech stimuli that is more cognitively demanding (e.g., in background noise) leads to greater cortical tracking (Yasmin et al., 2023). Similarly, we expected untrustworthy talkers to require additional cognitive demands, leading to stronger cortical speech tracking.

## Methods

### Participants

We recruited 20 right-handed, native English-speaking young adults (age range = 18 – 31, *M* = 23.95, *SD* = 4.20) with self-reported normal hearing. Eleven participants self-identified as men and nine as women. This sample size was chosen based on similar studies of cortical speech tracking (e.g., Di Liberto et al., 2015, 2018) providing sufficient data for stable estimation of TRFs (Di Liberto & Lalor, 2017; Mesik & Wojtczak, 2023). All participants provided informed consent before starting the experiment and received a €20 voucher for their time. All procedures were approved by the School of Computer Science and Statistics Research Ethics Board at Trinity College Dublin, The University of Dublin.

### Stimuli and Procedure

Participants completed a brief demographics questionnaire and the Propensity to Trust Scale (Evans & Revelle, 2008), which measures self-perceived trait-level trustworthiness and trust in others. Participants then completed the main experiment which took part in two rounds of two stages each. The first stage was a trust building game with gnome characters wherein participants tried to win as many gold coins as possible. The second stage involved listening to stories told by these same gnome characters. In each round, participants interacted with four different gnome characters. EEG data were recorded from 64 electrodes using a BioSemi ActiveThree system at a sampling rate of 512 Hz. Auditory stimuli were presented at 48 kHz through Sennheiser HD 650 headphones. The experimental task was programmed and run in MATLAB using functions from Psychtoolbox-3 (Brainard, 1997).

#### Trust-Building Stage

Participants were told they would play an investment game with gnome characters (**Figure 1A**). On each trial, they saw a coloured gnome icon and heard one of five possible investment rules (e.g., “Number of coins invested will be doubled”; see **Table 1**). Based on that rule, they chose how many gold coins to invest by pressing the corresponding number key (1–10; the “0” key = 10). They then received a return in coins. On some trials, the return matched the gnome’s stated rule; on others, it was reduced according to that gnome’s lie distribution.

**Table 1.** Investment Rules and Lie Distributions for Gnome Characters.

| **Investment Rules** |  |  |  |  |  |  |
|---|---|---|---|---|---|---|
| Number of coins invested will be doubled |  |  |  |  |  |  |
| Odd number: coins will be doubled, even number: receive 5 additional coins |  |  |  |  |  |  |
| Odd number: receive 5 additional coins, even number: coins will be doubled |  |  |  |  |  |  |
| Less than 5: coins will be doubled, 5 or more: coins will be multiplied by 1.5 |  |  |  |  |  |  |
| Less than 5: coins will be multiplied by 1.5, 5 or more: coins will be doubled |  |  |  |  |  |  |
| **Lie Distributions** |  |  |  |  |  |  |
| **Gnome** | **Frequency** | **Magnitude** | **Number of Lie Trials** | **Mean Return on Lie Trials** | **Expected Return per Trial** | **Mean Trust Rating (SE)** |
| 1 | Low | Low | 4 (20%) | 50% | 90% | 5.24 (0.27) |
| 2 | Low | Mid | 4 (20%) | 35% | 87% | 4.95 (0.36) |
| 3 | Low | High | 4 (20%) | 20% | 84% | 4.57 (0.25) |
| 4 | Mid | Low | 10 (50%) | 50% | 75% | 3.86 (0.35) |
| 5 | Mid | High | 10 (50%) | 20% | 60% | 2.48 (0.32) |
| 6 | High | Low | 16 (80%) | 50% | 60% | 2.77 (0.40) |
| 7 | High | Mid | 16 (80%) | 35% | 48% | 1.86 (0.28) |
| 8 | High | High | 16 (80%) | 20% | 36% | 1.33 (0.14) |

Gnome characters varied in terms of the frequency and magnitude of lies. For example, a gnome that was low in lie frequency but high in lie magnitude would lie on 20% of trials. On these lie trials, the return on investment would average 20% of the promised value (**Table 1** and **Figure 1** include further details on the lie distributions across gnomes). To differentiate the gnomes across lie distributions, we used eight different colours and eight different voices. Which colour and which voice was assigned to which lie distribution, as well as the corresponding order, was randomised for all participants. By utilising gnomes as opposed to human faces or more varied characters, and by randomising the voice-colour pairing across participants, we reduced the risk of trust bias stemming from variables unrelated to the trust game experience. After playing the game with four gnomes, participants then rated the perceived trustworthiness of each one on a scale from 1 – not at all to 6 – extremely using the corresponding buttons on the keyboard. After these trust ratings, participants moved on to the *story-listening stage* of the experiment.

#### 2.2.2 Story-Listening Stage

During the second phase of the experiment, participants were presented with a 5-minute story told by each gnome (**Figure 1B**). The same gnome colour and voice from the first stage were paired together, representing the same character from the first stage. The trustworthiness was fully established during the game stage. The trustworthiness/reputation was expected to transfer to the story-listening stage, where there was no additional deception or feedback. The story told by each gnome was randomised across participants. In each story, the gnome talked about a time where they were accused of doing something wrong and defended themselves from the accusation. Participants were reminded of their winnings from the *trust-building stage* before listening to each story and asked to rate whether or not they felt the gnome was innocent after each story. After listening to the four stories corresponding to the four gnomes they interacted with in the *trust-building stage*, the whole process repeated a second time (i.e., both stages with the four remaining gnomes).

### EEG data preprocessing

EEG data were pre-processed and analysed using custom MATLAB (MathWorks) scripts built starting from publicly available resources from the Cognition and Natural Sensory Processing (CNSP) initiative (https://cnspworkshop.net/). All data have been organised according to the Continuous Neural Data (CND) structure (Di Liberto et al., 2024), which is tailored to experiments involving continuous speech. All data will be shared in CND format upon publication. EEG data were band-pass filtered between 0.5 and 8 Hz, to include both the Delta and Theta EEG bands, which have both been shown to be central to the cortical speech tracking phenomenon (Chalas et al., 2023; Etard & Reichenbach, 2019). We used zero-phase shift 4^th^ order Butterworth filters and then down sampled to 64 Hz. Smooth spline interpolation was used to replace any bad EEG channels (more than three standard deviations away from the mean). EEG data were re-referenced to the average of all electrodes. As part of our secondary analyses, we also filtered the EEG data between 8 and 12 Hz (Alpha band) and pre-processed it in the same manner as above. Alpha power was computed by taking the squared value of the Hilbert envelope of the EEG signal for each channel and each sample. The values were then averaged across all channels.

### Stimulus Features

The primary EEG data analysis in this study aimed to estimate the neural encoding of speech at both the acoustic and linguistic levels. To that end, we extracted the amplitude envelope of each story, the derivative of the amplitude envelope and word onset times (**Figure 1B**). These features are widely used in TRF studies and reliably capture variance in the neural response during continuous speech perception (Brodbeck et al., 2018; Crosse et al., 2016; Lalor & Foxe, 2010).The amplitude envelope was computed by taking the absolute value of the Hilbert transform for each audio sample. The envelope derivative was calculated by taking the difference in the envelope magnitude between subsequent samples. Both features were then downsampled to 64 Hz to match the downsampled EEG. Word onset times were derived using the Montreal Forced Aligner (McAuliffe et al., 2017) which provided time-aligned lexical transcripts for each story. The resulting temporal alignments were manually inspected and corrected using Praat Software v.6.4.27 (Boersma, 2001) to ensure alignment accuracy, then converted to CSV format for further processing. Onset times were then converted to sample indices, and binary arrays were created for each story such that samples corresponding to word onsets were assigned a value of 1 and all other samples set to 0.

For secondary analyses, we also computed lexical surprisal and entropy values biased by different trust prompts to investigate how word predictability is affected by trustworthiness. Next word probability was estimated using the large language model Llama-2 (Touvron et al., 2023) with two different trust-related system prompts – “this speaker lies all the time” (Lie prompt) or “this speaker always tells the truth” (Truth prompt). Lexical surprisal is a measure of how unexpected a word is given the preceding context and was calculated as the negative logarithm of the extracted probability (Heilbron et al., 2022; Slaats & Martin, 2025). Lexical entropy measures the level of uncertainty the model has for next-word predictions given the context up to that point and was calculated as the weighted sum over the surprisal values for all possible next words (Goldstein et al., 2022; Slaats & Martin, 2025). This resulted in two surprisal and entropy values for each word of each story (i.e., one for each prompt). These values were then time-aligned to the word onsets and inserted into the corresponding feature arrays at the appropriate sample indices.

### EEG Analysis

EEG data were analysed using a temporal response function (TRF) approach (**Figure 1B**). TRFs estimate the linear mapping between a given sensory stimulus and the corresponding EEG responses (Ding et al., 2014; Lalor et al., 2009). TRFs were computed using a lagged linear regression approach that estimates a temporal filter for each EEG channel, capturing how the neural response at a given time point can be predicted from the preceding stimulus by considering a range of time lags. Here, we considered latencies between -100 ms to 600 ms (i.e., 0-600ms reflect neural reactions to the stimulus and include the TRF components typically associated with the stimulus features considered here; -100–0 enables the observation of possible neural activity preceding the stimulus, which would reflect anticipatory neural mechanisms).

For each participant, stories were separated into *High* and *Low* trust conditions based on a median split of the trust ratings at the end of the *Trust-Building Stage*. Separate forward TRF models were fit for each participant and trust condition to predict the EEG signal using a leave-one-out cross-validation procedure across trials to control for overfitting. Forward TRFs were fit using a ridge regularisation, with the regularisation parameter (λ) optimised through an exhaustive search over a logarithmic range from 0.01 to 100 within each training fold. The optimal λ was defined as the value yielding the highest prediction correlation (Pearson’s r) between the predicted and observed EEG signals (Crosse et al., 2016). EEG prediction correlations are an indication of the amount of EEG variance explained by a given feature or set of features.

## Results

The core hypothesis tested in our analysis was that the trust established in the *trust-building stage* influences the cortical tracking of speech in the *story-listening stage*. The first step was to verify numerically that the perceived trustworthiness was a reflection of the gnome lie distributions. Using a repeated measures ANOVA, we found that trust ratings varied across the lie distribution (*F*(7, 133) = 45.63, *p* < .001; **Figure 1A**). Specifically, all conditions resulted in significantly different trust ratings from one another (pairwise Wilcoxon signed-rank tests, FDR corrected, all *p* < .005), except among those with Low lie frequency (gnomes 1, 2, and 3 in **Table 1**). We also used repeated measures ANOVAs to verify that the trust ratings were not significantly different across the character voices (*F*(7, 133) = 0.84, *p* = .556) or colours (*F*(7, 133) = 0.39, *p* = .910), indicating that perceived trustworthiness was due to our manipulation and not to acoustic or visual biases (See supplementary materials for trust ratings broken down by voice and colour).

The next step was to test the main hypothesis that the established trustworthiness (*trust-building stage*) influenced the cortical tracking of speech (*story-listening stage*). Multivariate forward TRF models fit on High and Low trust gnomes separately (based on participant trust ratings; henceforth *High* and *Low* conditions respectively) were compared. Higher EEG prediction correlation values were measured for Low than High trust conditions (paired *t*-test, *t*(19) = 2.32, *p* = .032, *Cohen’s d* = 0.52; **Figure 2A**), reflecting a stronger low-frequency cortical speech tracking for less trustworthy talkers. We did not find any statistically significant differences when comparing individual electrodes across conditions, likely due to high inter-individual variability in the spatial distribution of responses across the scalp, which may be reduced by averaging across channels (FDR-corrected paired *t*-tests comparing EEG prediction correlations between conditions at each channel: all *p* > .05).

**Figure 2.**
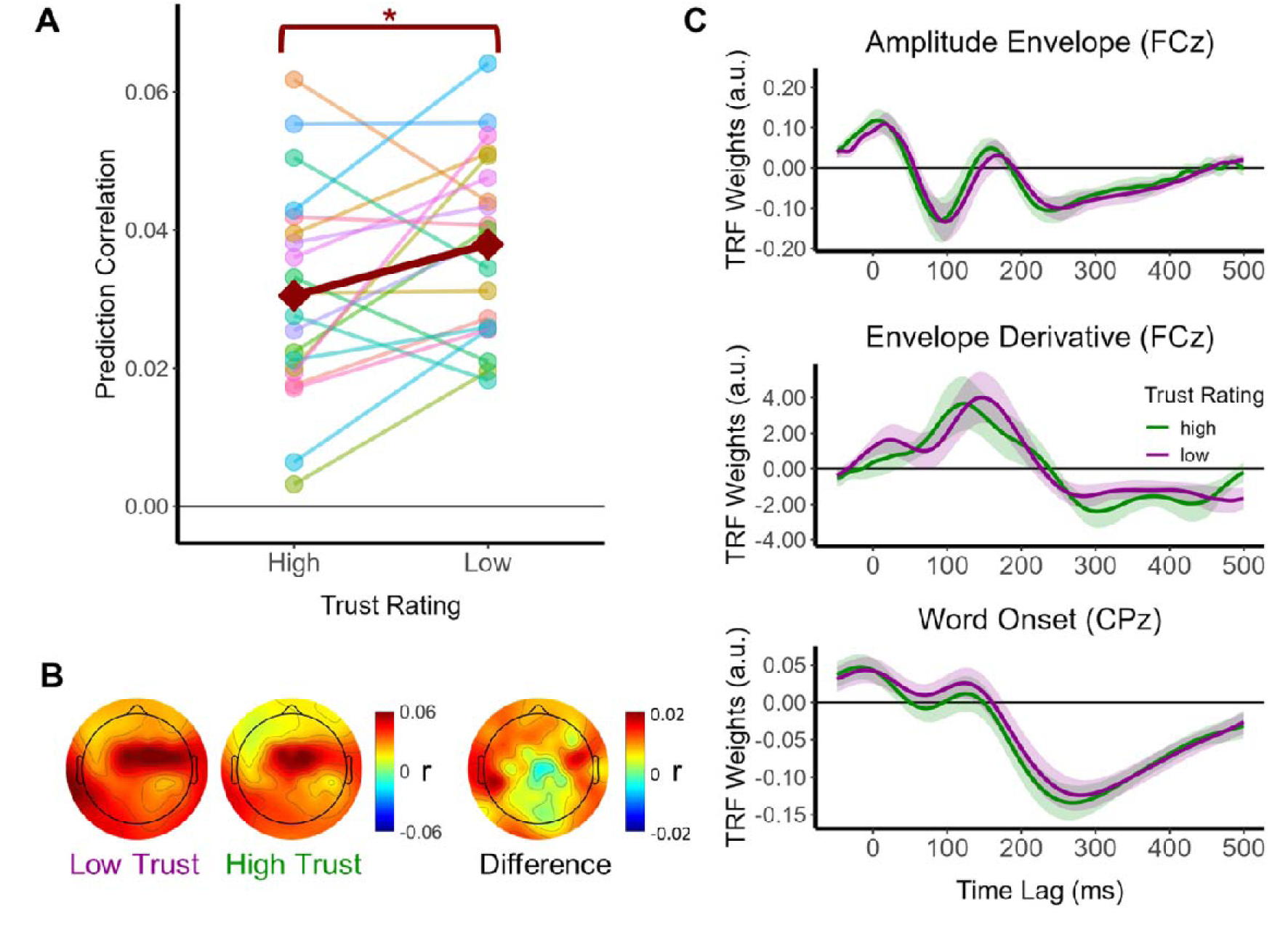
Distrust enhances the low-frequency cortical tracking of speech. **(A)** EEG prediction correlations values (average across all channels) grouped into High and Low trust conditions for each participant. The dark-red line represents the mean across all participants. Stronger EEG prediction correlations were measured when participants listened to untrustworthy individuals (* p < .05). **(B)** Distribution of EEG prediction correlations across the scalp for High and Low trust conditions, and their difference (Low minus High). Although prediction accuracy was significantly greater in the low-trust than the high-trust condition when averaged across electrodes, no individual electrode showed a statistically significant difference after correcting for multiple comparisons. **(C)** TRF weights for the amplitude envelope, envelope derivative, and word onset features for High trust (green) and Low trust (purple) conditions. No statistically significant differences were found between High and Low trust for any of the features at any time lags.

We then explored the possibility that trust influences how cortical signals represent speech. To that end, we tested if *High* and *Low* conditions corresponded with different TRF shapes for pre-selected channels. We looked at three central electrodes (Fronto-central: FCz, Central: Cz, and Centro-parietal: CPz). We found no statistically significant differences (FDR-corrected paired *t*-tests comparing TRF weights at each time lag all *p* > .05, **Figure 2C**), indicating no differences in peak timing or peak magnitude. Altogether, these results are in line with the view that trustworthiness influences the magnitude of the cortical speech tracking, without significantly altering the spatio-temporal dynamics of the underlying neural representation.

Further analyses were carried out to see what factors may be driving this trust effect in the prediction correlation. First, we assessed whether the effect of trust changed throughout a given story. To that end, we split each story in two and re-calculated the EEG prediction correlation on the first and second halves separately. The main effect of half was not significant (*F*(1, 76) = 0.07, *p* = .794, η*_g_^2^* < 0.01), indicating that the average prediction correlation across trust conditions did not vary between story halves. In line with the previous result in **Figure 2**, we found a main effect of Condition (repeated measures factorial ANOVA: *F*(1, 76) = 5.53, *p* = .021, η*_g_^2^* = 0.07) wherein the prediction correlations were greater for the Low condition (**Figure 3A**, see Supplementary Materials for individual participant trajectories). To further characterise the effect of condition, we ran paired t-tests for each story half and found a statistically significant difference between Low and High conditions in the first half of the story (paired t-test: *t*(19) = -3.67, *p* = .003, *Cohen’s d* = -0.82; FDR corrected), and not the second (paired t-test: *t*(19) = -0.84, *p* = .431, *Cohen’s d* = - 0.18; FDR corrected). The interaction between condition and half, however, was not significant (*F*(1, 76) = 2.09, *p* = .153, η*_g_ ^2^* = 0.03). Then we investigated if the increased prediction correlations in delta and theta bands correspond to differences in EEG alpha power, which has been associated with attention. Interestingly, there were no significant main effects or interaction between story half and condition (all *p* > .05), indicating that alpha power did not significantly vary across story halves and trust conditions.

**Figure 3.**
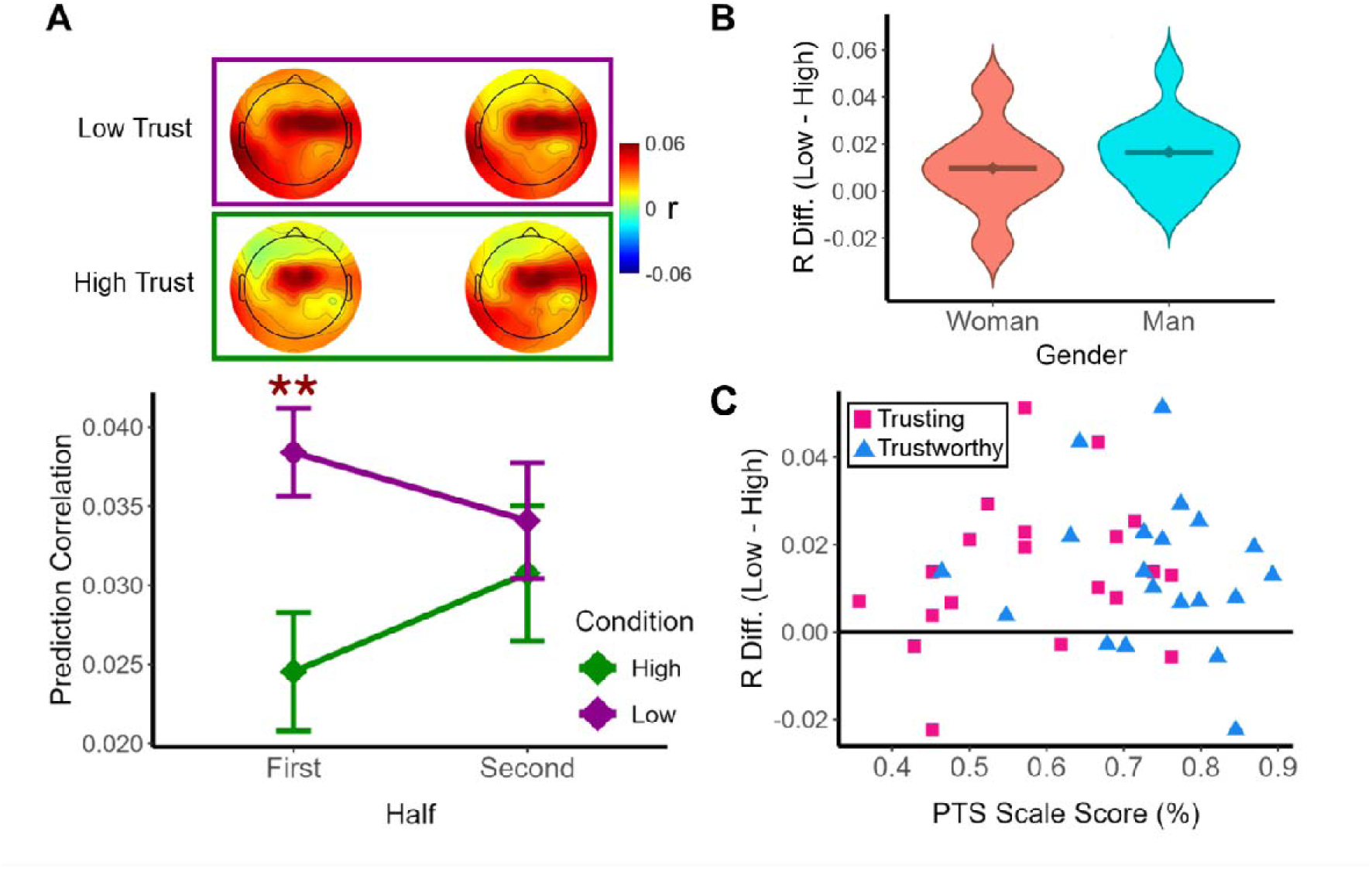
Previously established trustworthiness influenced cortical speech tracking only during the first half of story-listening. **(A)** EEG prediction correlations were re-calculated for the first and second halves of each story separately. Cortical speech tracking was different between Low and High trust conditions during the first half of the story but not during the second half. **(B)** We tested if the difference in EEG prediction correlation between trust conditions was related to gender. No statistically significant effect was found (p = .416). **(C)** We explore the possibility that the influence of trust on cortical speech tracking was related to individual propensity to trust. An indication of the individual propensity to trust was provided by the PTS scores, which included two metrics: how Trusting (pink squares) and how Trustworthy (blue triangles) a participant is. No statistically significant correlation was found between the change in EEG prediction correlation (Low minus High trust condition) and the PTS scores (Trusting: r = .215, p = .362; Trustworthy r = -.137, p = .564).

Finally, analyses were carried out to explore potential influences of demographic variables, such as gender or personality traits, on the effect of trust on the cortical tracking of speech. The EEG prediction correlations from the first half of the story, where the effect was stronger, were compared across men and women, finding no statistically significant difference (independent samples *t*-test, *t*(19) = - 0.84, *p* = .416, *Cohen’s d* = 0.19). Next, we tested for the possibility that the influence of trust on cortical speech tracking is correlated with the individual scale scores from the Propensity to Trust Scale (PTS), which provide numerical indices for trusting and how trustworthy a participant perceives themselves to be. We found no statistically significant correlation between the difference in EEG prediction correlation between Low and High trust conditions and how trusting someone is (*r* = .215, *p* = .362; **Figure 3C**), nor with how trustworthy they are (*r* = -.137, *p* = .564).

### Impact of Trustworthiness Biases on LLM next-word prediction

LLMs allow us to numerically assess how trustworthiness affects next-word prediction by using system level prompts. The First, we investigated whether the entropy and surprisal values derived from Llama-2 (Touvron et al., 2023) using the trust-based prompts are sensitive to trust. The entropy and surprisal values were significantly different between the two prompts for all stories. For surprisal, the Lie prompt resulted in significantly lower surprisal values than the Truth prompt across all eight stories (all *p* < .001). For entropy, we found the opposite – the Lie prompt resulted in significantly greater entropy values than the Truth prompt for all eight stories (all *p* < .001). Figure 4A shows the mean surprisal and entropy values for all stories. Although the values differed significantly across the two prompts for both entropy and surprisal, the correlations between the prompts were very high, especially for surprisal. For entropy, the Pearson correlation between the Lie prompt and the Truth prompt ranged from *r* = .659 to *r* = .957 across stories (mean *r* = .788). For surprisal, the Pearson correlations ranged from *r* = .947 to *r* = .993 (mean *r* = .963).

**Figure 4.**
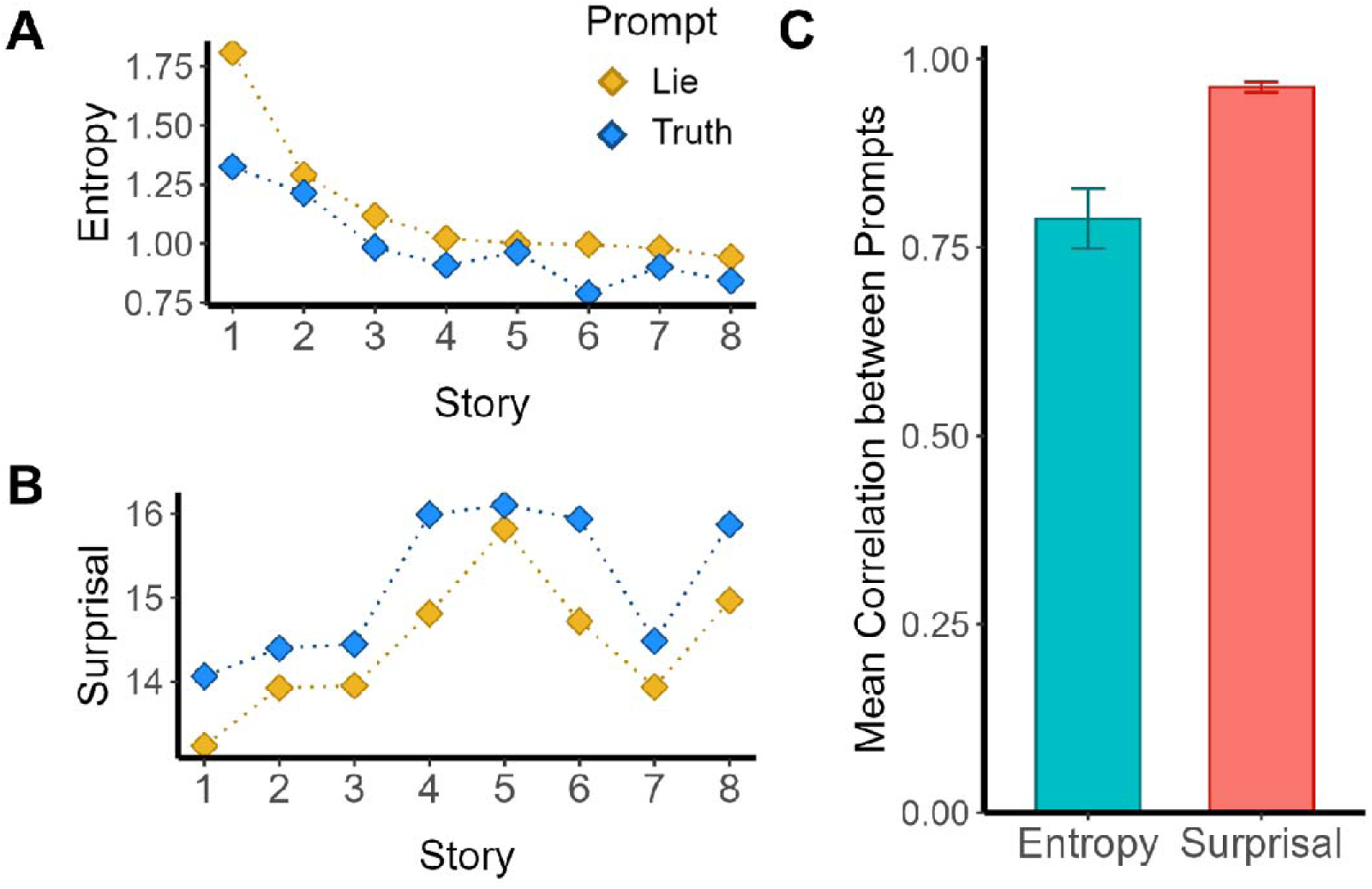
Word prediction from Llama-2 is sensitive to trust, but this is not reflected in neural tracking. **(A)** Entropy values (mean across all words) derived using Llama-2 are reported for each story when including a trustworthiness bias in the prompt. Entropy values were consistently higher when Llama-2 prompt included an indication that the story was told by a liar (i.e., Lie prompt) as opposed with a trustworthy individual (i.e., Truth prompt) (p < .001 for all stories). This indicates that distrust corresponds to an increased uncertainty in the lexical prediction. **(B)** Surprisal values were consistently lower for the Lie prompt than the Truth prompt (p < .001 for all stories). A lower surprisal is compatible with an increased entropy, as a more uncertain prediction tends also means less commitment to that prediction, hence less surprisal. **(C)** In addition to investigating the mean entropy and surprisal magnitude, we investigated if the trustworthiness bias influenced their temporal dynamics. We found strong correlations between Truth and Lie prompts ranging from .659 to .957 for entropy and from .947 to .993 for surprisal. Altogether, these results indicate that the trustworthiness bias primarily influences the mean magnitude of the entropy and surprise, and not their temporal dynamics.

Since TRFs would be sensitive to changes such as the ones in temporal dynamics (Figure 4C) but not to changes in the overall magnitude (Figure 4A,B), this points at next-word entropy as a candidate element of speech processing that might be particularly influenced by trustworthiness, which we think might be a useful starting point for future hypothesis testing. For example, we can investigate whether any differences between the prompts relate to TRF model performance (i.e., does the Truth prompt lead to better prediction correlations for the High trust condition than the Lie prompt and vice versa). We can also test if specific points in the story that are more sensitive to trust (based on differences between the prompts) also show greater differences in prediction correlation between High and Low trust conditions.

We ran additional multivariate TRF models on the *Low* and *High* conditions using the same features as before (amplitude envelope, envelope derivative, and word onset) and adding one of 1) entropy values from the Lie prompt, 2) entropy values from the Truth prompt. We then used a repeated measures factorial ANOVA to test if the prediction correlations were significantly different across conditions and prompts. We hypothesised that if Llama-2 was accurately modelling how trust impacts word prediction in the brain, we would find a significant interaction where the prediction correlation would be greater for congruent prompts (e.g., Lie prompt with Low trust) than incongruent prompts. As with the previous TRF analyses, we found a main effect of trust condition (*F*(1, 19) = 6.60, *p* = .019, η*_g_ ^2^* = 0.08) wherein the Low trust condition resulted in greater prediction correlations than the High trust condition. There was no significant main effect of LLM prompt (*F*(1, 19) = 1.05, *p* = .318, η*_g_ ^2^* < 0.01) or interaction (*F*(1, 19) = 0.47, *p* = .504, η*_g_ ^2^* < 0.01). This indicates that the different prompts are not significantly impacting model performance, with both prompts instead leading to higher prediction correlations for Low trust than for High.

Next, we wanted to investigate whether the differences between Low and High trust were related to the semantic content of the stories. To do this, stories were broken up into individual sentences. Any sentences of less than 6 words were combined with the sentence before or after (depending on which made more contextual sense). First, we took the predicted EEG from the models including amplitude envelope, envelope derivative and word onset and split it into individual sentences starting 100ms before first word onset and 600ms after the last word offset (to match the time lags from the TRF analysis). We then did the same for the actual EEG and found the prediction correlation between the two. To account for variance across subjects, prediction correlations were normalised by dividing by the subject’s mean across conditions. We then took the mean prediction correlation for each sentence across participants who heard it in the Low trust condition and those who heard it in the High trust condition. From here we took the difference in the mean prediction correlation between Low and High trust for each sentence from each story. The mean difference between the Lie and Truth prompt for each sentence was also computed for both surprisal and entropy. The sentences with greater differences between prompts should represent those that are more semantically related to trust. We looked at the correlation between the prediction correlation difference and the prompt differences for surprisal and entropy. These correlations were not significant across all stories (surprisal: *r* = .069, *p* = .163; entropy: *r* = -.041, *p* = .406). Altogether, these results indicate that the trustworthiness bias from the prompts influences the mean magnitude of the entropy and surprisal but does not mimic the temporal dynamics of trust in the human brain.

## Discussion

In the present study, we introduce a novel experimental framework for investigating how perceived trustworthiness impacts the neural processing of continuous speech. Trust was established during an initial behavioural stage using an investment game with fictional characters, and its impact was assessed during a subsequent story-listening stage. found that distrust increased cortical speech tracking, especially during the first half of the stories. Notably, this influence was specific to tracking strength: We found no evidence for changes in the representation of speech and language features in the neural signals. Together, these findings identify cortical speech tracking as a sensitive neural metric through which trust exerts its influence, as well as demonstrating a framework to study how this influence unfolds over time.

These results align with prior work showing that contextual factors can modulate the neural processing of continuous speech, even when the acoustic input is held constant (Yeshurun et al., 2017). Importantly, the effects of trust in this experiment are independent of prior biases associated with voice or appearance of the character. The experimental paradigm was explicitly designed to minimise and counter-balance such confounds. Both speech acoustics (Belin et al., 2017; Maltezou-Papastylianou et al., 2025) and visual features (Sofer et al., 2015; Willis & Todorov, 2006) are known to influence the perceived trustworthiness. To reduce visual biases, we used fictional characters (gnomes) rather than human faces, varying only colour across characters. Crucially, the pairing between character colour, voice, story, and trust-building stage trustworthiness was fully randomised across participants. As further confirmed by our numerical analyses, the aggregate results therefore reflect the influence of perceived trustworthiness itself. At the same time, this framework could be readily adapted in future studies to isolate and quantify the neural impact of specific trust-related biases.

Our findings indicate that trust modulates neural processing especially during the first half of each story. One possible explanation is that listeners dynamically revise their perceived trustworthiness of the speaker as the story unfolds. Alternatively, the prior trustworthiness of the character may remain stable, while its influence on story trust judgements fades over time (King-Casas et al., 2005; Mende-Siedlecki, 2018). In this view, the accumulation of linguistic context may progressively outweigh prior beliefs, rendering perception less susceptible to initial trust biases. Disentangling these possibilities will require future studies with more targeted manipulations of trust and narrative information.

The observed higher EEG-measured cortical speech tracking in the Low trust condition could arise from multiple underlying mechanisms. An intuitive explanation is that distrust enhances the alignment between neural activity and certain speech or language features. One possible perspective for this could come from Communication Accommodation Theory, which proposes that interlocutors adapt their communication behaviours to reduce social distance and increase convergence towards attractive and likeable communication partners (Gallois et al., 2005; Giles et al., 1973). Although this framework concerns alignment between communication partners rather than brain to speech alignment, it does raise the possibility that perceived trustworthiness could influence the degree to which listeners align with a communicative partner’s speech. From this perspective, we would predict greater cortical tracking for more trustworthy speakers. However, in line with our original hypothesis, low trust may increase cognitive load due to the additional uncertainty about the speaker’s intentions and reliability. When listeners perceive a speaker as unreliable, they may engage in enhanced monitoring of the incoming message to detect any inconsistencies or deception, increasing cognitive load and potentially enhancing the encoding of speech information. For example, distrust may increase the allocation of attentional resources (Daronnat et al., 2021; Fitzhugh et al., 2025) which would in turn could enhance EEG prediction correlations (Har-shai Yahav & Zion Golumbic, 2021; Yasmin et al., 2023). Here, we explored this possibility by analysing EEG alpha power. Interestingly, we found no differences between Low and High trust conditions.

Another possibility is that perceived trustworthiness influences lexical prediction mechanisms, leading to changes in EEG signal-to-noise ratio or in the alignment between LLM-features and EEG. To probe this idea, we injected trustworthiness biases into text prompts for the popular LLM Llama-2 (**Figure 4**). This manipulation revealed that distrust increased overall prediction uncertainty without altering word-by-word temporal relationships, a pattern consistent with a change in signal-to-noise ration rather than representational structure. While this analysis provides a useful intuition regarding how statistical predictions may be modulated by trust, it should be interpreted with caution, given that contemporary LLMs only partly capture the statistical and mechanistic properties of human language processing. For example, while LLMs exhibit human like effects on some reasoning tasks (Lampinen et al., 2024), even showing correlations between model embeddings and brain responses to language across time (Goldstein et al., 2022, 2025), their internal representations and output behaviours are ultimately driven by statistical pattern learning rather than lived experience or socio cognitive states. As a result, there are fundamental differences in how external context, such as speaker trust, is integrated. Additionally, system level prompts in LLMs are applied across the full context window, applying contextual constraints in a global, static manner (Kamruzzaman & Kim, 2025; Neumann et al., 2025), meaning they do not reflect the dynamic nature of trust we found in our TRF results. Nevertheless, we believe that utilising prompting in LLMs may provide a useful avenue for testing hypotheses related to different socio-cognitive states (e.g., familiarity between interlocutors, child-directed speech), especially by making use of different LLMs or prompt formulations.

Although the discussion so far has focussed on trust as the primary variable of interest, there are also scenarios in which trust may act as an important covariate that must be controlled rather than studied directly. This consideration is increasingly relevant as speech neurophysiology research shifts toward more naturalistic listening and interactive paradigms. For example, several studies use brief audiobooks, podcasts, or other single talker narratives (e.g., Broderick et al., 2018; Crosse et al., 2021; Lalor & Foxe, 2010; Panela et al., 2024). More recently, studies also incorporate dialogue listening (Ip et al., 2025) and even interactive dialogues between participants (Ishii et al., 2025; Chalehchaleh et al., in prep). Unlike traditional experiments involving isolated syllables or nonsense words, trust is intrinsic to real-world speech communication and even plays a role in non-interactive listening contexts, such as the consumption of informational or persuasive material. The framework introduced here provides tools for quantifying and accounting for trust-related effects, thereby enabling investigations of questions that are difficult or impossible to address behaviourally alone, such as how trust evolves over time during continuous speech listening, and how new information reshapes its influence on perception.

In conclusion, we show that perceived trustworthiness, a fundamental component of human communication, influences cortical speech tracking during the listening of continuous narratives. By combining an investment-based trust game with neurophysiological measurements during naturalistic speech listening, this study extends the trust literature into the domain of continuous speech perception. More broadly, it contributes to a growing body of work demonstrating that social and contextual factors shape speech perception at the neural level, thereby bridging traditionally distinct research domains. Further research is needed to identify the specific neural mechanisms underlying the changes in cortical speech tracking and to determine whether effects depend on different dimensions of trustworthiness, such as truthfulness, intent, or competence.

## Data Availability

Data will be made publicly available in CND format through the Open Science Framework (OSF) repository upon publication.

## Author Contributions

Jaimy Hannah: Conceptualization; Data curation; Methodology; Formal analysis; Writing—Original draft.

Giovanni M. Di Liberto: Conceptualization; Supervision; Funding acquisition; Writing—Review & editing.

## Acknowledgements

This research was conducted with the financial support of Research Ireland under Grant Agreement No 20/SP/8955 at the ADAPT Centre at Trinity College Dublin. ADAPT, the Research Ireland Centre for AI-Driven Digital Content Technology, is funded through the Research Ireland Centres Programme. The authors thank the Fidelity Center for Applied Technologies for their support, helpful discussions, and feedback throughout the project. The authors also thank Prof. Benajmin R. Cowan, Dr. Sophie Leonard, and Dr. Orla Cooney for regular discussion on the progress of this study, including reflections on experimental design and results.

## Notes

### Competing Interest Statement

The authors have declared no competing interest.

### Summary of Updates

Minor changes throughout the text. Figure 1 has been revised.

