## supplementalInfo for "Trust Modulates Speech Entrainment: Enhanced Cortical Tracking for Low-Trust Speakers"

**Supplementary Materials**

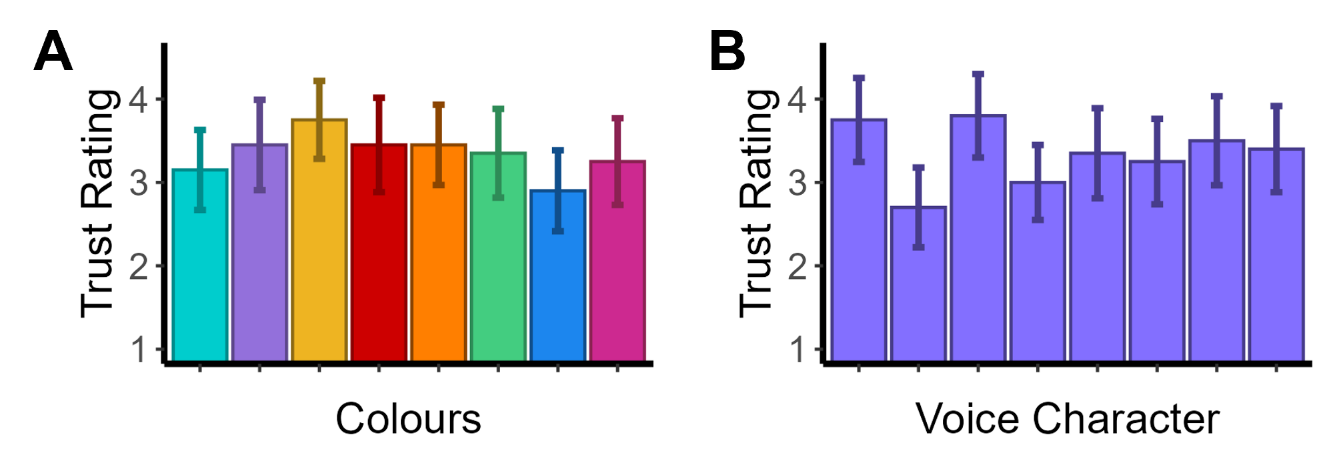

***Figure S1. Trust ratings by gnome colour and voice. (A)*** *Trust ratings did not significantly differ across colours (F(7, 133) = 0.39, p = .910), with mean trust values ranging from 2.90 (dark blue) to 3.75 (yellow) on a 1-7 scale. Error bars represent standard error.* ***(B)*** *Trust ratings did not significantly differ across voices (F(7, 133) = 0.84, p = .556), with mean trust values ranging from 2.70 to 3.80 on a 1-7 scale. Error bars represent standard error.*

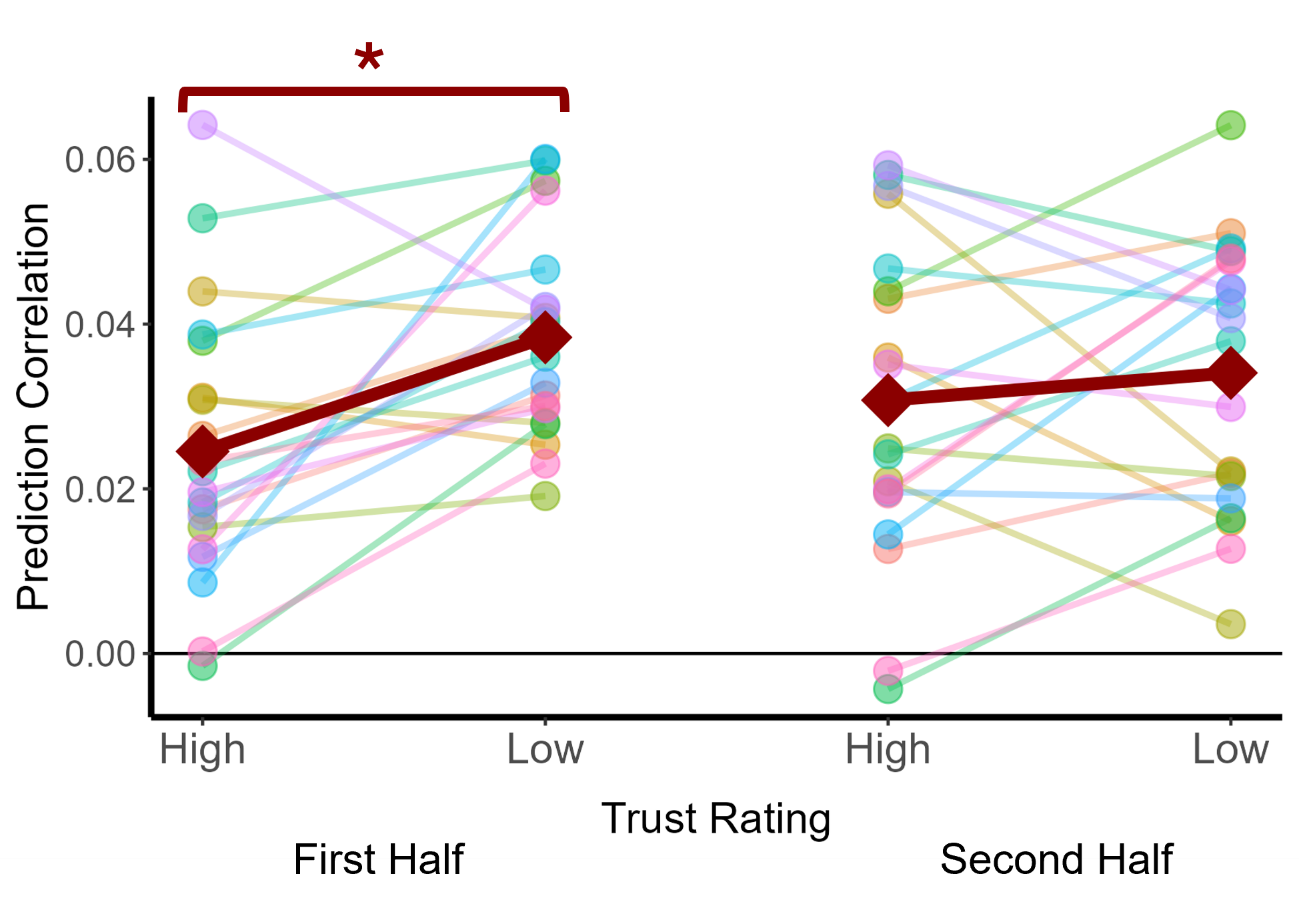

***Figure S2. Prediction correlation values by Trust Rating and Story Half.*** *During the first half of the story, prediction correlations were significantly lower for High trust ratings than for Low trust ratings. During the second half of the story, the difference between trust conditions was not significant. The red diamonds indicate the mean prediction correlation across participants in each condition. The lighter coloured lines each represent one participant.*
